# Combined aerobic and resistance exercise training alters the spatial transcriptome of skeletal muscle in young adults

**DOI:** 10.1101/2024.06.07.597971

**Authors:** Michael J. Stec, Zachary A. Graham, Qi Su, Christina Adler, Min Ni, Valerie Le Rouzic, David R. Golann, Patrick J. Ferrara, Gabor Halasz, Mark W. Sleeman, Kaleen M. Lavin, Timothy J. Broderick, Marcas M. Bamman

## Abstract

Chronic exercise training substantially improves skeletal muscle function and performance. The repeated demands and stressors of each exercise bout drive coordinated molecular adaptations within multiple cell types in muscle tissue, leading to enhanced neuromuscular recruitment and contractile function, stem cell activation, myofiber hypertrophy, mitochondrial biogenesis, and angiogenesis, among others. To comprehensively profile molecular changes induced by combined resistance and endurance exercise training, we employed spatial transcriptomics coupled with immunofluorescence and computational approaches to resolve effects on myofiber and mononuclear cell populations in human muscle. By computationally identifying fast and slow myofibers using immunofluorescence data of spatially sequenced tissue sections, we identified fiber type-specific, exercise-induced gene expression changes that correlated with muscle functional improvements. Additionally, spatial transcriptome profiling and integration of human muscle single cell RNAseq data identified an exercise-induced shift in interstitial cell populations coincident with angiogenesis. Overall, these data provide a unique spatial molecular profiling resource for exploring muscle adaptations to exercise, and provide a pipeline and rationale for future studies in human muscle.

## Introduction

Exercise is a highly dynamic and potent stimulus that results in large-scale changes to the molecular, cellular and tissue landscape. Acute exercise, regardless of the mode (e.g., continuous endurance, resistance, etc.), creates a unique stress that requires highly-coordinated intra– and inter-cellular communication to maintain and optimize performance while also regulating cellular processes responsible for restoring energy sources, removing damaged proteins and cellular debris, and returning the cell to homeostasis^1,2^. Repeated and progressive exercise training results in a number of beneficial outcomes that span cognitive, psychological, physical, and cellular domains that greatly improve health and reduce/prevent the risks of chronic disease onset^3–5^, ultimately leading to improved quality of life and healthspan.

Skeletal muscle is the predominant actor during exercise. It is the most abundant tissue in a normal, healthy individual, making up approximately 40-45% of total body mass. The specific adaptations of skeletal muscle to progressive exercise training are specific to the exercise prescription. In broad terms, continuous aerobic exercise at moderate to moderate-high intensities is associated with improving oxidative metabolic function via mitochondrial biogenesis and capillary angiogenesis, while resistance training induces in muscle hypertrophy via net protein synthesis over time. Some of the mechanisms behind these processes have been reported extensively^2,4^. Our molecular understanding of exercise-induced adaptations of skeletal muscle has exploded in recent years, leveraging advances in high-resolution profiling technologies such as RNA sequencing and concomitant advances in bioinformatics and computational biology.

Skeletal muscle is a complex tissue containing, in addition to primary myofibers, numerous non-myofiber cell types and associated structures (e.g., neural, vascular, connective, and adipose tissues along with immune cells, stem cells, etc.). Traditional approaches to muscle transcriptomics have mostly relied on bulk sequencing of RNA isolated from whole tissue^6–9^ or from single cell/nuclei isolated from the muscle environment^10–12^. While these data have been invaluable, they are incomplete as they lack important architectural information about the local cellular environment and origin of the RNA as the tissue adapts to exercise training. Spatial transcriptomics overcomes this limitation using histological sections that are either labeled using in situ hybridization techniques or high-throughput spatially-defined barcodes (e.g., 10X Genomics Visium pipelines) followed by sequencing to allow for transcriptomic assessment within the native tissue environment, without the need for disruptive cell/nuclei isolation procedures which can substantially impact gene expression^13–15^. Isolating regions, co-imaging with serial sections immunostained for histological features, and deconvoluting data using advanced computational and statistical models may allow for a deeper understanding of changes in the muscle transcriptome with exercise training that are unique to the many cell types. Recent reports have demonstrated the power of spatial transcriptomics in pre-clinical models of muscle damage^16,17^ and denervation^18^, and we have recently used it to identify skeletal muscle spatial profiles in models of muscular dystrophy^19,20^. However, to our knowledge, no study has investigated how highly supervised exercise training alters the muscle spatial transcriptome of human participants. Here, we performed spatial transcriptomics with multiplexed immunofluorescence histology, coupled with downstream computational pipelines to isolate myofiber type sub-populations and integration of single cell RNAseq data to define mononuclear cell populations. We found that that 12 weeks of supervised combinatorial exercise training results in myofiber type-specific changes in the transcriptome as well as interstitial cell population shifts that are associated with biological and functional changes in muscle, highlighting the utility of this analysis approach for providing a comprehensive profile of molecular adaptations to exercise within the native tissue environment.

## Results

### Spatial gene profiling of human muscle biopsies identifies exercise-responsive cell clusters

To determine the effects of exercise training on the skeletal muscle spatial transcriptome, healthy young male (n=3) and female (n=2) adults (Table 1) were recruited for supervised combinatorial aerobic and resistance exercise training for 12 weeks (exercise protocol detailed in STAR Methods). Importantly, the combined exercise training included multiple exercises targeting the knee extensors (e.g., vastus lateralis), including treadmill running, cycle ergometry, back squat, and loaded knee extension. Resting vastus lateralis muscle biopsies were collected in the fasted state before training at week 0 (sedentary, Sed.) and after (exercised, Ex.) the 12 week exercise program. A portion of the biopsy (∼50 mg) was immediately processed and frozen for subsequent cryosectioning, immunostaining, and spatial gene expression profiling using the Visium CytAssist workflow (Figure 1A). After muscle sections were immunostained and imaged for myofiber type, transcriptomic probes were hybridized to the tissue and then transferred to Visium slides containing 55µm diameter spots with spatially barcoded capture probes for downstream sequencing. On average, 690 probe spots were analyzed per sample, with a median depth of 29,972 reads sequenced per spot (Table S1). In order to create a joint embedding space, data integration was performed per participant to enable joint analysis of all sequenced samples. The integrated data showed separation between sedentary compared to exercised samples (Figure 1B), as well as clear separation into 9 clusters that correspond to distinct cell types (Figure 1C), which were annotated based on their canonical gene expression profiles (Figure 1D, Table S2).

**Figure 1.**
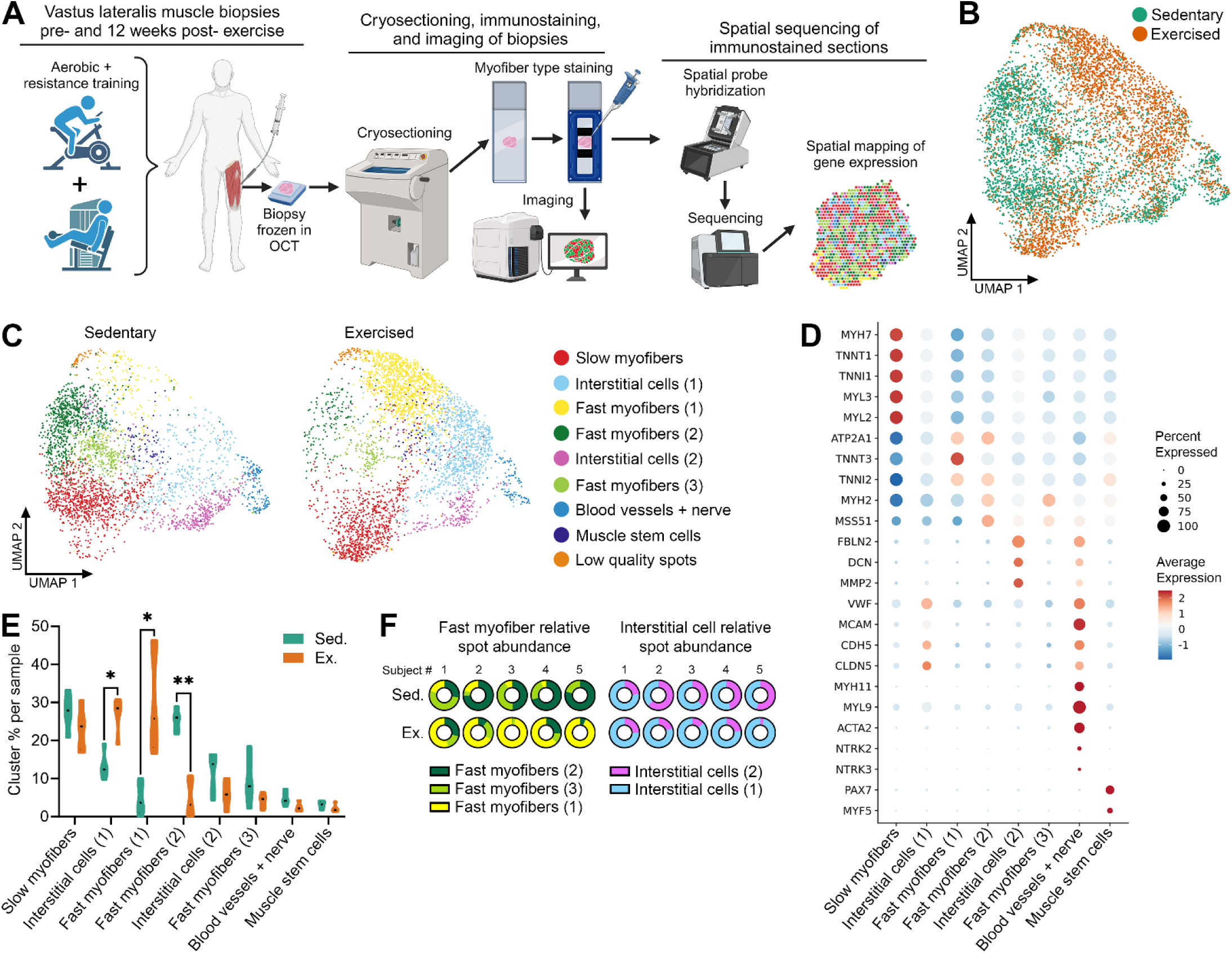
Spatial transcriptomics identifies unique cell clusters in human muscle biopsy sections, including exercise-responsive clusters. (A) Schematic outlining procedures for exercise training, muscle biopsy collection and processing, and spatial transcriptomics analysis. (B) UMAP clustering of probe spots from all samples, highlighting differences between sedentary and exercised samples. (C) Unbiased clustering of spots from all samples, identifying 9 unique clusters annotated by canonical gene expression profiles. (D) Dot plot showing the expression of canonical genes used to annotate each cluster. (E) Percentage of cluster spots per total spots within each sample, comparing spot abundance in sedentary vs. exercised samples; * p < 0.05, **p < 0.01. (F) Proportion of Fast myofibers (1), (2), and (3) clusters among the total Fast myofibers population within each sample and proportion of Interstitial cells (1) and (2) among the total Interstitial cells population within each sample, comparing distribution in sedentary vs. exercised samples.

**Table 1.** Subject Characteristics.

|  | Males (n = 3) | Females (n = 2) |
| --- | --- | --- |
| Age (years) | 19.8 ± 1.2 | 22.2 ± 3.4 |
| Body fat (%) | 25.8 ± 8.3 | 37.8 ± 1.7 |
| Thigh lean mass (kg) | 13.8 ± 1.2 | 9.1 ± 0.5 |
| VO<sub>2</sub>peak (mL/kg/min) | 35.0 ± 5.4 | 27.9 ± 2.2 |

Data are presented as mean ± SD.

While all identified clusters contained some degree of expression of myofiber-specific genes due to the 55µm diameter probe spot overlapping areas with myofibers, several clusters were clearly enriched for myofiber gene expression. For example, we identified a cluster with spots enriched for slow myosin heavy chain gene expression (*MYH7*), as well as other slow contractile genes (e.g., *TNNT1*, *MYL2*, *MYL3*, *ATP2A2*), and thus, annotated this cluster as Slow myofibers. In contrast, several fast myofiber clusters [annotated as Fast myofibers (1), (2), and (3)] were identified based on their high expression of fast myofiber genes, including *MYH2*, *TNNI2*, and *TNNT3*. In addition to myofiber clusters, we also identified two populations that were enriched for genes expressed by muscle interstitial cells [Interstitial cells clusters (1) and (2)], including extracellular matrix (ECM)-producing cells (*FBLN2*, *DCN*, *MMP2*, and multiple collagen genes) and endothelial cells (*VWF*, *MCAM*, *CDH5*, *CLDN5*). Finally, spatial profiling identified spots annotated as blood vessels + nerve based on smooth muscle marker expression (*MYH11*, *MYL9*, *ACTA2*) and a minor proportion of spots containing nerve markers (*NTRK2*, *NTRK3*). Further, spatial resolution was sufficient to detect a cluster annotated as muscle stem cells based on the expression of canonical markers including *PAX7* and *MYF5*. A low abundance cluster (<2% of all spots) did not display a remarkable gene expression profile, and it was found that spots of this cluster localized to the edges of tissue sections that contained swollen/damaged myofibers consistent with freeze artifact (Figure S1), and thus this cluster was denoted as “low quality spots”.

To further characterize the multiple subtypes of fast myofiber and interstitial cell clusters, we plotted their relative abundance in both sedentary and post exercise training samples. Interestingly, among the fast myofiber clusters, clusters (2) and (3) were predominantly enriched in sedentary muscle, while fast myofibers cluster (1) was significantly enriched (6.6-fold greater abundance) in exercise trained muscle (Figures 1E-F). Similarly, interstitial cells cluster (2) was found mainly in sedentary samples, while interstitial cells cluster (1) was significantly enriched in exercise trained samples (2-fold greater abundance).

Overall, unbiased spatial transcriptomics profiling of human muscle clearly identified unique clusters with expression profiles of different myofiber types, interstitial cells, blood vessels, and muscle stem cells. Interestingly, a robust transcriptomic response to 12 weeks of exercise training in fast myofibers and interstitial cell populations led to the identification of novel clusters enriched in exercise trained muscle. These data support the utility of spatial transcriptomics in not only identifying localized areas of gene expression associated with different cell types in human muscle tissue, but also the ability to detect cell type-dependent changes in gene expression following exercise training.

### Distinct tissue localization of spatial clusters supports cell type identification

While the gene expression signatures of each cluster provided strong evidence for the cell types within clusters, a benefit of spatial profiling coupled with multiplexed immunofluorescence directly on the sequenced tissue sections is that cluster spots can be mapped to histological features to further confirm cell type identity. From a global perspective, there was an increase in fast myofibers (1) and interstitial cells (1) cluster spots present throughout exercised samples compared to sedentary samples, however, there was no clear pattern of cluster dispersion throughout the tissues for any of the clusters (Figure 2A), consistent with the vastus lateralis having a heterogenous mosaic pattern distribution of myofiber types and other cell types in healthy young individuals. When visualizing spot localization of individual cell clusters, spot overlay with corresponding immunofluorescence images revealed distinct overlap of spots with cell type and tissue architecture that supported their annotation (Figure 2B). The typical cross-sectional diameter of human myofibers in the vastus lateralis of young adults ranges 50-90µm (i.e. cross-sectional area ∼2000-6400µm^2^) while cross-sectional areas of other muscle-resident cells (e.g., fibroblasts, macrophages, endothelial cells, stem cells, pericytes, etc.) are much smaller. Thus, 55µm diameter probe spots typically overlayed parts of multiple myofibers and other cell types. However, the slow myofiber cluster spots overlayed predominantly myofibers containing type I (slow) myosin heavy chain protein, and most fast myofibers spots overlayed mainly myofibers containing type IIa (fast) myosin heavy chain. Aside from immunostaining for myofiber type, laminin and nuclear staining revealed that ‘blood vessels + nerve’ spots overlayed structures with the morphology of blood vessels, and ‘interstitial cell’ cluster spots could be found at least partially overlapping interstitial spaces between myofibers. Overall, these data indicate that gene expression signatures of multiple clusters matched their cell type identification by immunofluorescence, thereby confirming accurate annotation of spatial clusters.

**Figure 2.**
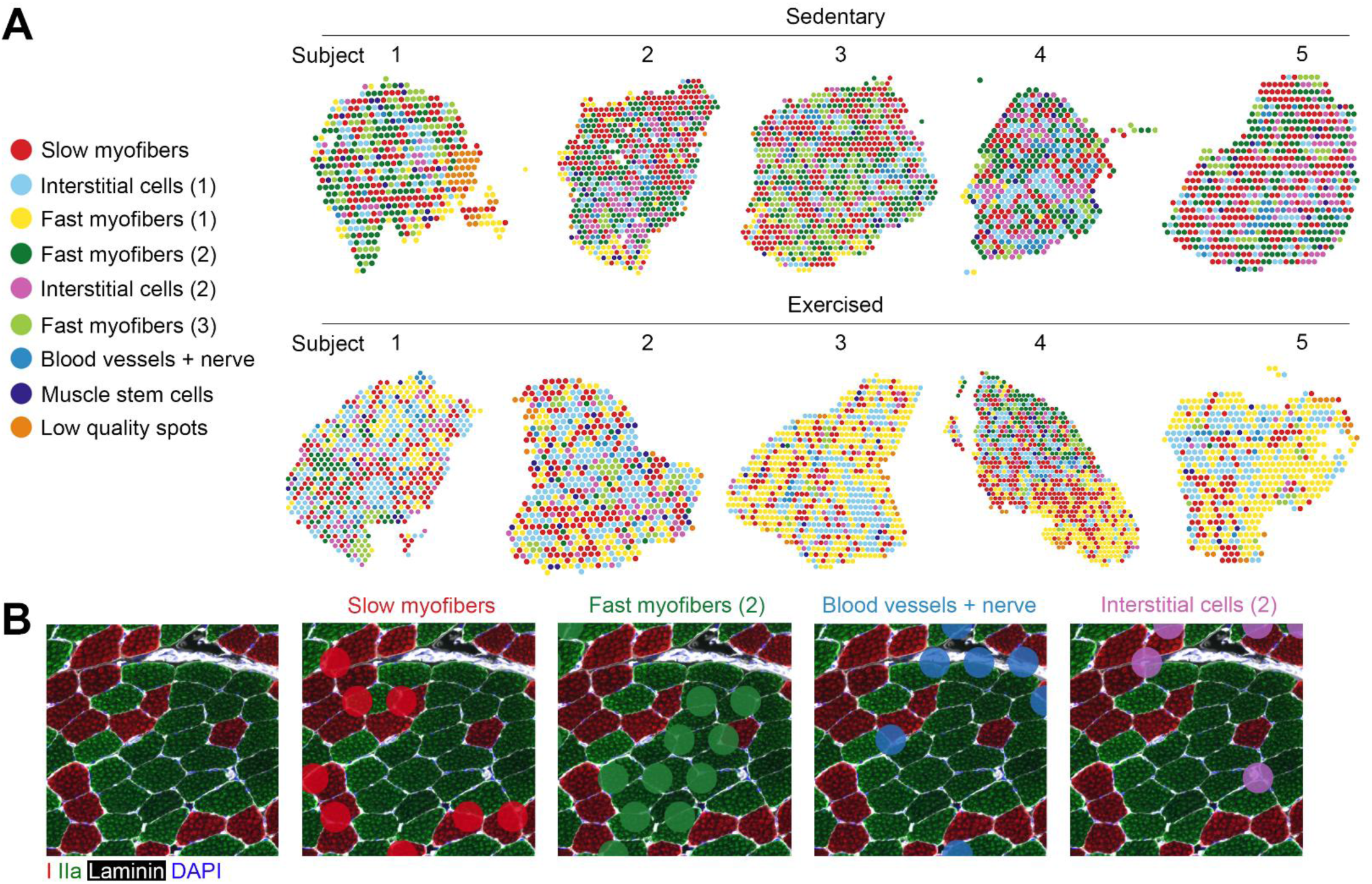
Localization of cell clusters identified by unbiased gene signatures corresponds with predicted cell types. (A) Distribution of each unique cluster throughout entire biopsy area for all samples before and after exercise training. (B) Overlay of probe spots with immunofluorescence histological markers for annotated clusters; spot size = 55µm.

### Refining myofiber spot classification via spatial immunofluorescence signal yields more accurate myofiber type identification

To identify spots specifically overlaying only slow or fast myofibers, we extracted myosin heavy chain type I and type IIa immunohistochemistry (IHC) intensity data from spot areas of the sequenced samples to distinguish areas containing only a single myofiber type. Staining intensity data from red (type I myosin) and green (type IIa myosin) channels within all spots were extracted, and cutoffs for each channel were set at the second turning point within the intensity distribution (Figure 3A). Spots with staining intensity above the cutoff for a single myofiber type and below the cutoff for the other myofiber type were considered to contain almost exclusively a single myofiber type, while spots that were either above or below the cutoff for both myofiber types were not considered to be a pure myofiber population (Figure 3B). From this IHC-based analysis, newly classified slow and fast myofiber spots aligned more accurately with their respective myofiber type compared to previously defined clusters (Figure 3C), and these spots contained significantly greater intensity of staining for their respective myosin heavy chain, while having a lower intensity of staining for the other myosin isoform (Figure 3D). Additionally, when comparing myofiber spots within IHC staining intensity thresholds compared to myofiber spots not meeting IHC threshold criteria, gene expression profiling revealed that IHC-defined myofiber spots had a more robust myofiber type-specific signature. For example, differential gene expression analysis comparing IHC-defined fast myofiber spots with fast myofiber spots outside the IHC intensity threshold revealed the IHC-defined fast myofiber cluster had enriched expression of fast contractile genes, including *MYH1*, *MYH2*, *MYLPF*, *TNNT3*, and *TNNI2*, and reduced expression of slow contractile genes, including *MYH7*, *MYL2*, *MYL3*, *TNNT1*, and *TNNI1* (Figure 3E). Similarly, the IHC-defined slow myofiber cluster displayed enriched slow myofiber contractile genes and reduced fast myofiber genes compared to slow myofiber spots outside the IHC intensity threshold (Figure 3E). Altogether, while unbiased clustering of gene expression signatures defined multiple slow and fast myofiber clusters, leveraging myosin heavy chain IHC staining intensity was a more accurate method to identify clusters containing more pure myofiber type populations.

**Figure 3.**
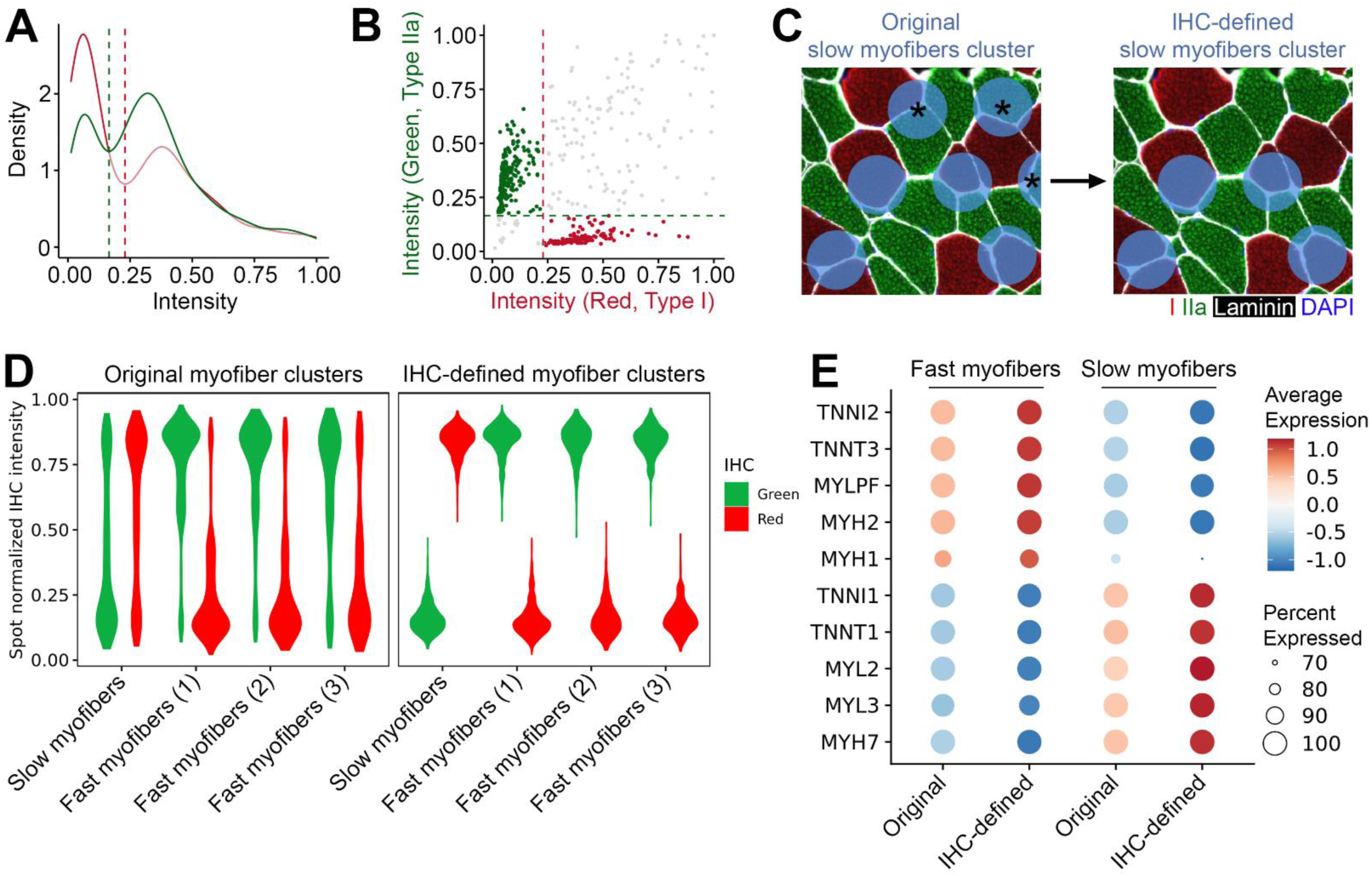
Extracting multiplexed immunohistochemistry (IHC) data from sequenced tissue sections improves detection of different myofiber type clusters. (A) Example of spot density distribution within a single sample based on computationally detected immunofluorescence intensity within each spot (red = type I myosin, green = type IIa myosin) (B) Cutoff parameters (dotted lines) used to isolate spots containing pure myofiber types, based on the second turning point within intensity distribution depicted in (A). Each dot represents a single probe spot within the sample, with red dots representing IHC-defined slow (type I) myofiber cluster spots, green dots representing fast (type IIa) myofiber cluster spots, and gray dots representing spots that did not meet myofiber type cutoff thresholds. (C) Left panel: unbiased detection of probe spots based off gene expression signatures annotated as slow myofibers, with asterisks highlighting several spots overlapping both slow myofibers (red stain) and fast myofibers (green stain). Right panel: improved identification of slow myofiber-specific spots using IHC-based analysis workflow, where the IHC-defined slow myofiber spots encompass mainly type I stained areas. (D) Violin plot showing the normalized IHC intensity of green (type IIa) and red (type I) fluorescent signal for spots within the originally defined myofiber type clusters from unbiased gene expression analysis (left) and from the IHC-defined myofiber type analysis (right). (E) Dot plot showing gene expression of canonical fast and slow myofiber markers, comparing expression between the originally defined myofiber type clusters to the IHC-defined myofiber type clusters.

### Exercise-induced gene expression profiles of fast and slow myofiber clusters are coincident with improved muscle function

To determine myofiber type-specific gene expression changes regulated by exercise training, differential gene expression analysis was performed on IHC-defined fast and slow myofiber clusters. Overall, 12 genes were significantly upregulated and 82 downregulated in the fast myofiber clusters, and 20 genes were upregulated and 96 downregulated in the slow myofiber clusters (Table S3). Gene ontology analysis revealed significant changes in Cellular Components pathways containing myofibril genes in fast myofibers, as well as sarcoplasmic and sarcoplasmic reticulum genes in both fast and slow myofibers (Figure 4A). Additionally, mitochondrial matrix and mitochondrial protein-containing complex pathways were uniquely altered by exercise in the slow myofiber cluster only. Genes upregulated in the fast myofiber cluster in exercise trained samples included *PLN* and *SLN*, whose protein products regulate sarcoendoplasmic reticulum calcium ATPase (SERCA) activity, as well as the myosin binding protein *MYBPH*, which has previously been found to be upregulated after resistance training^21^, and *ACTN3*, a fast-twitch α-actin that is known to be associated with strength and power (reviewed in^22^) (Figure 4B). Genes upregulated in the slow myofiber cluster in response to exercise included regulators of mitochondrial function (e.g., *SOD2*, *MT-ND2*) and angiogenesis (*VEGFB*) (Figure 4C).

**Figure 4.**
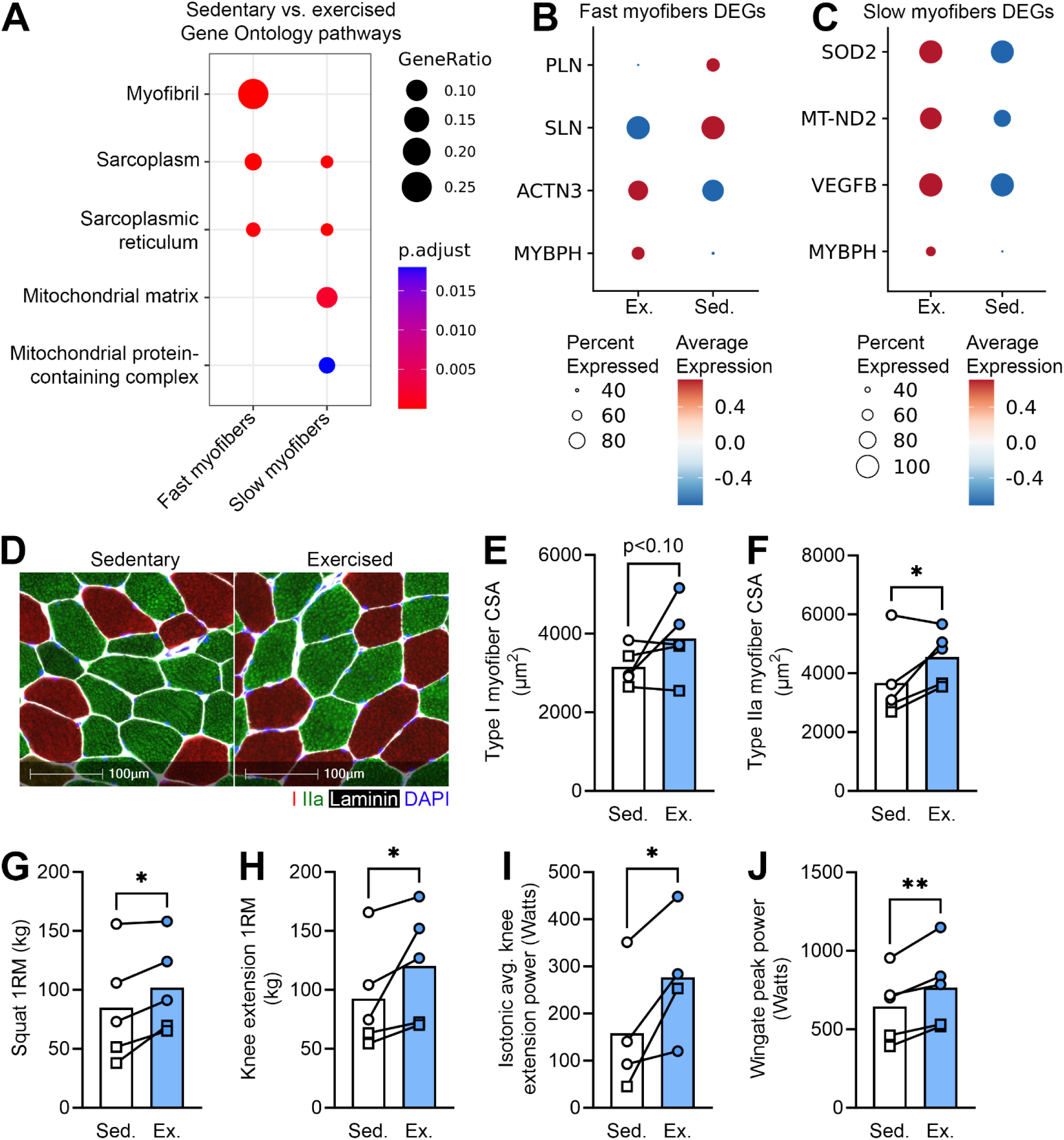
Exercise uniquely alters gene expression profiles of fast and slow myofiber clusters, with concomitant increases in muscle function. (A) Gene Ontology pathways (Cellular Components) differentially regulated between sedentary vs. exercised samples in IHC-defined fast and slow myofiber clusters. Gene ratio is the total number of differently expressed genes divided by the total number of genes in each gene set, and the adjusted p-value is based on gene set enrichment analysis. (B-C) Relative expression levels differentially expressed genes between sedentary and exercised samples in IHC-defined fast and slow myofiber clusters. (D-J) Exercise-induced gene expression changes within fast myofiber clusters are associated with morphological and functional changes consistent with fast myofiber adaptation to exercise, including increased type IIa myofiber cross-sectional area (CSA) (D-F), increased squat (G) and knee extension (H) one repetition maximum (1RM), increased isotonic knee extension power (I), and increased peak power assessed by Wingate power testing (J). Circles represent data points from male participants and squares represent data points from female participants. * p < 0.05, **p < 0.01

In concordance with beneficial gene expression changes in fast myofiber clusters, morphological and functional changes supported fast myofiber adaptations to exercise training. For example, there was a significant increase in type IIa myofiber size (and a trend towards increased type I myofiber size) from pre-to post-training (Figure 4D-F). Additionally, multiple functional parameters indicative of fast myofiber adaptation, including enhanced muscle force and power production, were increased from pre-to post-training (Figure 4G-J). Altogether, leveraging IHC intensity data to derive fast and slow myofiber clusters enabled identification of myofiber type-specific changes in gene expression in response to exercise training, indicating altered contractile gene expression in fast myofibers and mitochondrial and angiogenic gene expression in slow fibers, with fast myofiber gene expression changes coinciding with improved fast myofiber morphology and function.

### Differences in interstitial cell clusters are associated with enhanced capillary density following 12 weeks of exercise training

Aside from changes in gene expression in myofiber clusters between sedentary and exercise trained samples, training drove a clear shift in the populations of interstitial cell clusters, evidenced by a significant increase in interstitial cells (1) and decrease in interstitial cells (2) cluster spot abundance in exercise trained samples compared to sedentary samples (Figure 1E-F). To understand the change in the interstitial cell landscape driven by exercise, we leveraged human muscle single cell RNAseq data from Rubenstein et al.^10^ to computationally infer the cell types present in interstitial spaces in sedentary and exercised samples. This single cell dataset contains multiple cell types, including endothelial cells, fibroadipogenic progenitors (FAPs), smooth muscle cells, pericytes, and immune cells. We calculated gene module scores for these single cell populations and integrated these scores with each spatial transcriptomics cluster gain a better understanding of the cellular composition of each cluster (Figure S2). Confirming the cell type identity of spatial clusters, muscle stem cells cluster had the highest module score for ‘satellite cells’, while the blood vessels + nerve cluster had the highest module scores for populations including ‘smooth muscle cells’ and ‘pericytes’ from the Rubenstein et al.^10^ data (Figure S2E-G). Interestingly, spots within the interstitial cells (2) cluster, which was more abundant in sedentary samples, had significantly higher gene module scores for ‘FBN1+ FAP cells’ and ‘LUM+ FAP cells’ compared to interstitial cells (1) cluster (Figure 5A-B); whereas the interstitial cells (1) cluster, which was more abundant in exercise trained samples, had significantly higher gene module scores for ‘Endothelial cells’ and ‘Pericytes’ (Figure 5C-D).

**Figure 5.**
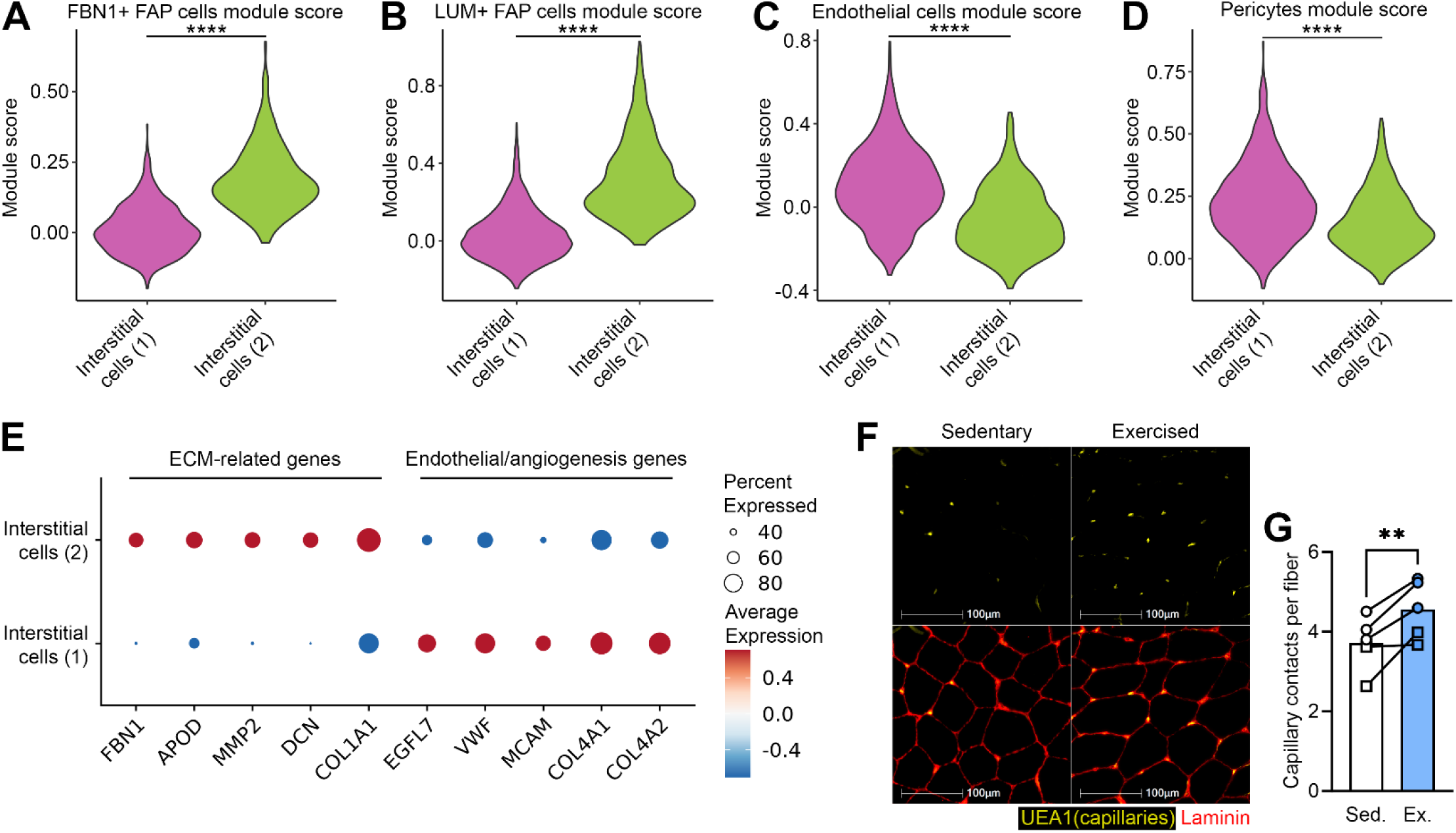
Spatial profiling reveals an altered cellular landscape of interstitial cell clusters following exercise, driven by a shift from fibroadipogenic to endothelial cell abundance. (A-D) Violin plots depicting integration of gene module scores for fibroadipogenic (FAP) cells (A-B) and endothelial and pericyte cells (C-D) taken from Rubenstein et al.^10^ single RNAseq cell datasets onto probe spots of Interstitial cells (1) and (2) clusters. (E) Dot plot showing differences in gene expression of ECM-related genes and endothelial/angiogenesis-related genes between Interstitial cells (1) and (2) clusters. (F) Immunostaining for UEA1 to detect capillaries, and quantification of the average number of capillary contacts per fiber in sedentary and exercised samples. Circles represent data points from male participants and squares represent data points from female participants. **p < 0.01

To complement single cell RNAseq integration that suggested a shift from FAP populations towards endothelial/pericyte populations with exercise training, we performed differential gene expression analysis between interstitial cells clusters (1) and (2). Gene ontology analysis of differentially expressed genes revealed that biological processes including ‘response to transforming growth factor beta’ and ‘extracellular structure organization’ were altered between interstitial cell clusters (Table S4), suggesting a potential enrichment for fibrogenic cells within one of the clusters. As predicted from single cell RNAseq integration analysis, interstitial cluster (2), which was more abundant in sedentary samples, had heightened expression of genes known to be expressed in mouse and human FAP cells and fibroblasts, including *FBN1*, *APOD*, *MMP2*, *DCN*, and *COL1A1* (Figure 5E)^10,23–25^. Additionally, gene ontology analysis identified the biological process, ‘regulation of angiogenesis’, as being differentially regulated between interstitial cell clusters (Table S4). Endothelial marker genes, as well as genes known to promote angiogenesis, including *EGFL7*, *VWF*, *MCAM*, *COL4A1*, and *COL4A2*, were significantly enriched in interstitial cluster (1), which was more abundant in exercise trained samples, again suggesting a potential increase in angiogenic processes and endothelial cell abundance in exercise trained samples (Figure 5E). In concordance with these observations, immunostaining of serial sections of these samples revealed a significantly increased number of capillary contacts per myofiber in exercise trained vs. sedentary samples (Figure 5F-G), supporting increased endothelial cell abundance within the interstitial cell population during training. Overall, these data demonstrate that despite using a relatively large spot diameter of 55µm for spatial gene detection, unbiased spatial clustering of spots, along with integration of human muscle single cell RNAseq data, can accurately predict shifts in muscle interstitial cell compositions following 12 weeks of exercise training.

## Discussion

Exercise training-induced changes in skeletal muscle have been well characterized at the whole body, isolated system, myofiber, and molecular levels^2,4^. The remarkable plasticity of skeletal muscle in its adaptability to resistance training, endurance training, or the two in combination, requires complex cellular responses. The data presented here are the first to describe exercise training-induced, spatially-resolved transcriptomic changes in human skeletal muscle after combined training, highlighting how cell-dependent molecular responses are associated with functional adaptations to progressive resistance and endurance training.

With the rapid advancement of transcriptomics technology, the ability to probe molecular responses to exercise has improved; however, the unique cellular composition and organization of skeletal muscle makes it difficult to generate a comprehensive transcriptome among all cell types. Skeletal muscle tissue is comprised primarily of multinucleated myofibers, which are not suitable for single cell RNAseq platforms due to their large size and multiple nuclei. While single myofibers as well as other cell types can be isolated from human muscle biopsies for sequencing, these disruptive procedures can result in substantial changes in gene expression, evidenced from mouse experiments where mechanical and enzymatic isolation of muscle mononuclear cells/nuclei (e.g., muscle stem cells, FAPs) were found to alter expression of up to ∼2,000 genes^13–15^. Additionally, single cell and single nucleus RNAseq technologies do not allow for spatial resolution, which is important in the context of skeletal muscle, as myofiber type patterning is altered in aging, disease^26,27^, and exercise training. Recently, spatial transcriptomics technology has been used to profile animal models of neuromuscular/musculoskeletal perturbations^16–20^. Here we show for the first time that this technology presents a clinically feasible and robust method for the molecular profiling of fresh frozen human muscle biopsies.

Previous high resolution single cell/nucleus sequencing and deconvolution of bulk muscle transcriptomics from human samples have detected approximately a dozen different cell types present in the muscle environment, varying by donor condition and clustering approach^10,12,24,25,28^. Here, unbiased clustering of spatial gene expression signatures identified 8 unique cell clusters, including two cluster groups that were uniquely modulated by chronic exercise training. In line with previous spatial transcriptomics studies in mice, we were able to identify multiple clusters of myofibers, including slow and fast myofiber types. Fast myofiber type transitioning is a hallmark of exercise training, with reduced abundance of hybrid fibers and loss of type IIx fibers seen with both endurance and resistance exercise^29^. While we were unable to distinguish a clear profile of IIx vs. IIa or hybrid myofiber clusters based on gene expression changes of these myosin heavy chain isoforms, unbiased clustering was able to identify three unique populations of fast myofibers, one of which was significantly enriched in exercised samples.

To further explore exercise-induced changes in both fast and slow myofiber cluster transcriptomes, we first established a computational method to identify probe spots containing only fast or slow myofibers. Unlike the clear spatially compartmentalized myofiber type distribution in mouse muscle, human vastus lateralis muscle biopsies from healthy young individuals contain a heterogenous distribution of myofiber types, whereby a 55µm probe spot can overlap areas containing multiple myofiber types, thus confounding gene expression data within that spot. To overcome this limitation, we extracted intensity data from multiplexed IHC staining for myosin heavy chain proteins (performed on the same tissue sections that were sequenced) and were able to define threshold cutoffs for identifying spots that were mostly pure fast or slow myofibers. This analysis significantly enhanced our resolution of myofiber types, as evident by a clear myosin heavy chain staining profile within IHC-defined spots, as well as a more pure gene expression profile of either fast or slow myofiber-related genes. After computationally defining fast and slow myofiber cluster spots, differential gene expression analysis between sedentary and exercised samples revealed myofiber type-specific changes in gene expression, with fast myofibers displaying changes in myofibrillar genes and slow myofiber displaying changes in mitochondrial genes. Among the genes altered following exercise training in the IHC-defined fast myofiber cluster, the top upregulated gene, *MYBPH*, has previously been noted to be more highly expressed in fast vs. slow myofibers^30^, and has also been shown to be among one of the most increased genes detected by bulk transcriptomics following resistance exercise training in young adults^21^. Additionally, *ACTN3*, which has previously been found to be exclusively expressed in fast myofibers^31^ and is a gene that is known to be associated with strength and power phenotypes in humans (reviewed in^22^), was among the top three upregulated genes in exercised vs. sedentary samples. In concordance with the presumably beneficial upregulation of these genes in the fast myofiber cluster in exercised samples, multiple parameters assessing fast myofiber function were improved following exercise, including increased fast myofiber size (type IIa myofiber cross-sectional area), increased muscle strength (squat and knee extension 1 repetition maximum), and increased muscle power (isotonic knee extension and Wingate peak power). These data provide proof of concept that spatial transcriptomics coupled with our computational approach for identifying spots containing single myofiber types can be used to accurately identify myofiber type-specific gene signatures that correspond with associated changes in function.

Aside from using spatial transcriptomics to detect exercise-induced changes in the abundant myofiber populations, we also had the resolution to detect other less abundant cell types, including clusters defined as Muscle stem cells, Interstitial cells, and Blood vessels + nerve. While overlaying probe spots on IHC images was able to validate that the Blood vessels + nerve cluster likely contained these cells/structures, we instead integrated human muscle single cell data from Rubenstein et al.^10^ to confirm the identity of other clusters that were unable to be visualized by only the laminin and nuclear staining performed on the sequenced samples. Integration of single cell gene module scores confirmed that the Muscle stem cells cluster had the highest module score for ‘satellite cells’, and that Interstitial cell clusters contained high module scores for single cell populations including FAPs, endothelial cells, and pericytes. Interestingly, we found that the Interstitial cells cluster (2) that was enriched in sedentary samples had a significantly higher module score for FAP populations, and higher gene expression for ECM-related genes and FAP markers compared to the Interstitial cells cluster (1) that was more abundant in exercised samples. On the other hand, the exercised-enriched Interstitial cells cluster (1) had high gene module scores for endothelial cells and pericytes and displayed heightened gene expression of angiogenic and endothelial genes compared to Interstitial cells (2) cluster. These gene module enrichment and gene expression data suggested a shift in interstitial cell populations from FAPs towards endothelial/pericyte cells with exercise in our spatial transcriptomics study, and intriguingly, Rubenstein et al.^10^ also found that after 12 weeks of resistance training, there was a significant increase in the endothelial/pericyte population defined by single cell RNAseq. As capillary density and capillary contacts per fiber have been shown to increase with both endurance and resistance training^32,33^, we postulated this computationally inferred shift towards endothelial/pericyte populations would manifest in increased capillarization in exercised samples. Indeed, when we immunostained serial sections for the detection of capillaries, we found significantly increased capillary contacts per myofiber in exercised vs. sedentary samples. Overall, these data demonstrate that spatial transcriptomics technology can be used to not only identify myofiber type-specific changes in gene expression driven by exercise, but that it can also be used to inform changes in composition of the interstitial milieu in response to exercise.

While our data provide a valuable resource of a skeletal muscle spatial transcriptome that highly correlates with functional and phenotypic outcomes of a combinatorial exercise intervention, these data are not without limitations. A major limitation of this study, as well as other published muscle spatial transcriptomics studies, is due to the 55µm spot diameter of the current 10X Genomics Visium pipeline that leads to spots overlapping with multiple cells, including myofibers of different types, as well as different mononuclear cell populations. We were able to partially mitigate this limitation by developing a computational approach to isolate spots that contained myofibers of a single type and by integrating human muscle single cell data to identify different mononuclear cell populations, though our resolution would be further improved by next generation spatial transcriptomics technologies with smaller probe spot sizes. Another limitation of this study is that there is heterogeneity between samples inherent of human participant muscle biopsies. A biopsy site may be repeated across time, but it is impossible to knowingly duplicate the exact biopsy site, and thus, repeat biopsies from a single subject may yield different fiber types and mononuclear cell populations. Despite this, the overall patterns of our data among these subjects as a whole are consistent with profiles of healthy young muscle, as well as known exercise-induced adaptations. Finally, a potential limitation of this study is that we studied the spatial transcriptome following chronic (12 weeks) exercise training as opposed to an acute exercise bout (e.g., minutes to hours after training). While our spatial dataset following chronic training was optimal for identifying gene expression changes associated with myofiber remodeling and for detecting shifts in interstitial cell populations driven by repeated exercise bouts, future studies examining the muscle spatial transcriptome after an acute bout of exercise may provide more robust changes in gene expression and identification of cell type-specific changes that drive adaptation to exercise.

In summary, we generated a spatial transcriptome of human skeletal muscle by leveraging immunostaining of sequenced tissue sections combined with a computational approach to enrich for myofiber types, as well as integration of human single cell transcriptome data for delineation of mononuclear cell types. Our data show that this pipeline yields accurate identification and localization of cell types as well as exercised-induced shifts in cell populations and cell type-dependent changes in gene expression that are associated with phenotypic and functional changes in muscle. Future studies can employ this spatial transcriptomics analysis approach in a variety of contexts, for example, in human muscle biopsies following acute exercise to detect cell type-dependent transcriptomic changes driving muscle adaptations, or in muscle biopsies from individuals with age-related or neuromuscular disease-driven myofiber type grouping to elucidate spatial queues regulating myofiber denervation/reinnervation.

## Acknowledgements

This study was supported by Department of Defense Office of Naval Research (N000141613159; TJB, MMB) and the Department of Veterans Affairs Rehabilitation Research and Development Service (IK2RX002781; ZAG). Illustrations were created with BioRender.com.

## Author Contributions

TJB and MMB designed the parent exercise trial. MJS, ZAG and MMB conceived and designed the research herein. MJS, QS, CA, VLR, DRG, and PJF completed experiments. MJS, ZAG, QS, and MMB analyzed and interpreted results. MJS and QS generated figures. MJS and ZAG wrote the manuscript. MJS, ZAG, QS, MN, PJF, GH, MWS, KML, TJB, and MMB edited and revised the manuscript.

## STAR Methods

### Resource Availability

#### Lead Contact

Further information and the datasets generated and analyzed for this study may be directed to and will be fulfilled by Lead Contact, Michael Stec.

#### Materials Availability

This study did not generate new unique reagents.

#### Data Availability

- Spatial transcriptomics data will be deposited in the Gene Expression Omnibus (GEO) managed by the National Center for Biotechnology Information.
- This paper does not report original code.
- Any additional information required to replicate the data reported in this paper is available from the lead contact upon request.

### Experimental Model and Study Participant Details

#### Participants

Biospecimens used in this study were obtained from a parent clinical trial comparing traditional combined exercise versus high-intensity combined exercise prescriptions (Department of Defense, Office of Naval Research: N000141613159, ClinicalTrials.gov ID: NCT0330923). We have previously described the recruitment, eligibility, screening and randomization strategies for these participants^7^. In brief, males and females aged 18-27 were recruited from the greater Birmingham, AL area. They were sedentary (no regular exercise history in the past 12 months), non-smokers, generally healthy and had a BMI < 30. The study was approved by the University of Alabama-Birmingham Institutional Review Board (Protocol: F160512012) and all participants provided informed consent to allow the use of their biospecimens for future research.

#### Exercise training

All samples used in this study were from participants (n = 5; 3 male, 2 female) that completed the ‘Traditional’ arm (TRAD) of the parent exercise dosing trial. All participants in the trial attended 3 supervised sessions per week for 12 weeks. Those in the TRAD arm completed a 30 minute bout of steady state exercise using either a treadmill or stationary cycle ergometer at 70% heart rate reserve followed immediately by ∼60 minutes of resistance training. Resistance training was comprised of 3 sets of 13 repetitions using a superset design that included lower body exercises (back squat, knee extension, heel raise) and upper body exercise (chest press, overhead press, seated row, lat pulldowns, triceps push-downs, and biceps curls), along with abdominal crunches. Participants were given 60-75 seconds rests between sets and the different exercises. Treadmill speed, cycling resistance, and weights were progressively increased as needed throughout the trial.

#### Dual-energy x-ray absorptiometry

Body composition was assessed via total body dual-energy x-ray absorptiometry (DXA) using a GE Lunar iDXA (GE Healthcare, Chicago, IL, USA). All scans were analyzed using GE Encore software v18 with primary variables of this report being total body fat and bilateral thigh lean mass.

#### Aerobic capacity testing (VO_2_peak)

VO_2_peak was determined using a continuous ramp protocol on a LODE Excalibur Sport cycle ergometer with sex-specific progression. For females, initial resistance was 50 W with an increase of 8.3 W/min for the first 9 min followed by a 15 W/min increase for the remainder of the test. For males, initial resistance was 75 W with an increase of 8.3 W/min for the first 9 min followed by increases of 25 W/min thereafter. Heart rate (HR) was measured continuously with a 3-lead ECG. Blood pressure was collected every other minute (all even minutes) while rating of perceived exertion (RPE) was collected every other minute starting 1 min after the start of the test (all odd minutes). The test was terminated when the participant reached volitional fatigue or was unable to maintain a cadence above 60 RPM. The main VO_2_peak achievement criteria were volitional fatigue and achieving a respiratory exchange ratio (RER) > 1.10.

#### Maximal anaerobic power testing

Anaerobic power was measured using the standardized 30 s Wingate cycle test on either a LODE Excalibur Sport cycle ergometer Monark cycle ergometer. Participants performed a 5 minute warm-up with no load prior to the test. Per standard protocol, males were tested at a resistance of 0.7 kiloponds per kilogram bodyweight and females were tested at a resistance of 0.67 kiloponds per kilogram.

#### Knee extensor strength and power testing

Peak unilateral maximum voluntary isometric contraction strength and dynamic power were evaluated on the dominant leg using a Biodex System 4 Pro dynamometer. Peak knee extension power was determined against a constant external resistance set at 40% of the maximal voluntary isometric contraction(MVIC) torque. Participants completed five consecutive, full range of motion unilateral repetitions as rapidly as possible during the concentric phase. Peak power was defined as the highest power output achieved across the five concentric repetitions while average power was defined as the average of the five power peaks.

#### One repetition maximum (1RM) squat testing

Prior to baseline testing, participants performed 3 familiarization sessions with the back squat exercise. For both baseline and post-training testing, participants performed low intensity warmups with progressively increasing weight. Successful 1RM attempts were followed by two minutes of rest, followed by subsequent attempts with increased weight until failure. 1RM was determined as the preceding weight that was successfully completed after two failed attempts at an increased weight.

#### Muscle biopsies

Muscle biopsies were collected during rest following an overnight fast before the start of training (sedentary) and at the end of training (exercised) using the Bergstrom technique from the vastus lateralis^34^. A portion of the biopsy was placed in a mixture of Optimal Cutting Temperature compound and tragacanth gum and subsequently frozen in liquid nitrogen cooled isopentane for histological analyses. Histological mounts were shipped to Regeneron overnight on dry ice for sectioning and spatial transcriptomics.

#### Tissue processing and immunofluorescence staining

Muscle biopsies were cryosectioned at 10µm thickness onto SuperFrost Plus charged slides and stored at –80°C until staining and processing via the Visium CytAssist pipeline. Muscle sections were stained according to the manufacturer’s protocol (10x Genomics, Rev D). In brief, samples were fixed with ice cold methanol, blocked, incubated with primary antibodies (Type I, DSHB Cat #BA-F8; Type IIa, DSHB Cat #SC-71; Laminin rabbit; Sigma Aldrich Cat #L9393), washed, incubated with secondary antibodies (mIgG2b 546, Invitrogen Cat #A21143; mIgG1 647, Invitrogen Cat #A21240; rIgG 488, Invitrogen Cat #A32731; Hoechst 33342, Invitrogen Cat #H3570), washed, immersed in 3x SSC buffer, and mounted with glycerol. Slides were immediately imaged using a Zeiss AxioScan.Z1.

For follow-up staining to detect muscle capillaries, serial sections were fixed with 4% paraformaldehyde (PFA) for 15 minutes, washed three times with PBS, and were then incubated in blocking solution containing 4% bovine serum albumin and 0.3% Triton X-100 in PBS for one hour. Sections were incubated with laminin antibody (Sigma Aldrich Cat #L9393) overnight in blocking solution, washed with PBS, and then incubated with anti-rabbit secondary (Invitrogen) and FITC-conjugated UEA-I (Sigma-Aldrich Cat #L9006) for one hour at room temperature. Sections were then washed with PBS, mounted in Fluoromount G (ThermoFisher Cat #00-4958-02), dried overnight, and imaged using a Zeiss AxioScan.Z1.

#### Visium spatial library construction and data processing

The 10x Genomics Visium CytAssist Spatial Gene Expression for FFPE platform was used for spatial transcriptomics experiments. Tissue sections mounted on standard glass slides were hybridized to the 10x Genomics human transcriptome probe set for CytAssist (v2) according to the manufacturer’s protocol (10x Genomics). The CytAssist machine was used to transfer hybridized probes from the tissue sections to Visium 6.5mm slides. Spatially-tagged cDNA libraries were built using the 10x Genomics Visium CytAssist Spatial Gene Expression for FFPE Kit. Sequencing was performed on an Illumina NovaSeq 6000 (Read 1 = 28, Read 2 = 50, Index 1 = 10, and Index 2 = 10). Alignment to the human reference genome GRCh38.93 was done using the Space Ranger 2.0.0 pipeline to derive a feature spot-barcode expression matrix (10x Genomics).

#### Visium data analysis

Data analysis was done in R (version 4.2), using the Seurat package (version 4.3). Spots with few gene counts or low counts of total number of UMIs were excluded from analysis. The cutoffs were: nFeature >= 500 and nCount >= 1000. Upon filtering, sedentary and exercised samples from the same donor were merged by using the Suerat merge function. Upon merging, datasets from each donor were normalized using SCTransform and then integrated (FindIntegrationAnchors). Hereafter, Principal Component Analysis was applied to the integrated data, and then top 25 principal components were used to generate a neighborhood graph. Last, data was clustered with resolution at 0.5. Cluster-specific enriched genes (Table S2) conserved between sedentary and exercise were identified by using FindConservedMarker with default setting. The top markers per cluster were based on Bonferroni-corrected p-values. Differentially expressed genes between sedentary and exercised samples were identified by using FindMarker with default setting.

#### Spots classification

Png package (version 0.1.8) was used to extract staining intensity data from both red and green channels of the IHC stained images. Based on the distribution of spot color intensity data from both red and green channels, the cutoffs for each channel were determined by locating the second turning point within the corresponding intensity distribution. This method was applied individually to each sample’s IHC stained image, to account for potential differences in brightness of each sample.

### Quantification and Statistical Analysis

Statistical analyses were performed using GraphPad Prism software (version 10.1.0). Paired student’s t-tests were used to compare the % of spots per sample for each cluster in sedentary vs. exercised samples, changes in myofiber CSA and muscle function in sedentary vs. exercised conditions, single cell gene module scores between interstitial cell populations, and the number of capillary contacts per fiber in sedentary vs. exercised samples. Data are reported as mean ± SEM, with significance at p < 0.05. Five samples per group were analyzed for each experiment.

## Supplemental Information Legends

**Figure S1.**
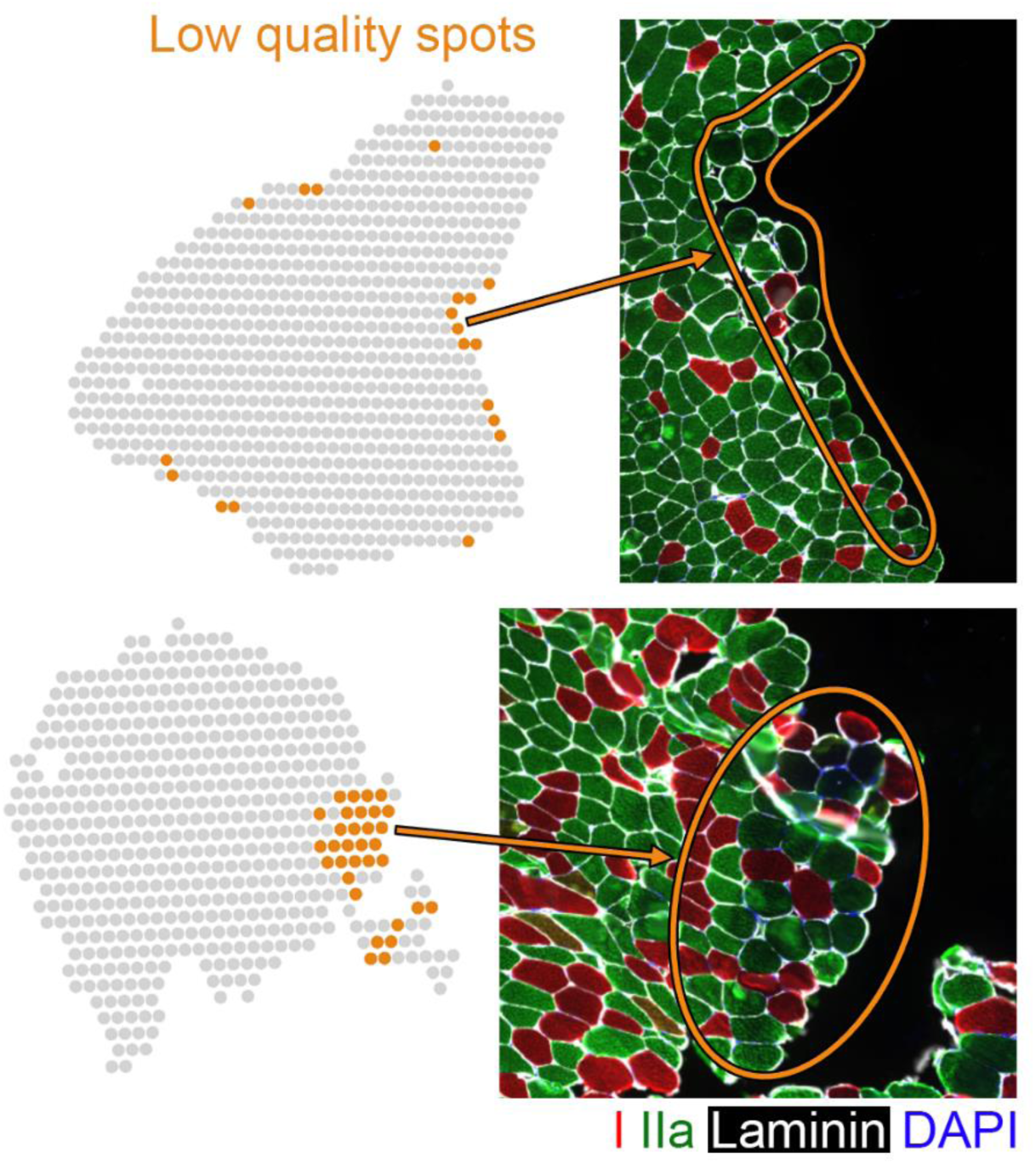
“Low quality spots” cluster typically associates with regions near tissue edges and contains swollen/damaged myofibers. Representative images depicting spatial distribution of low quality spots within entire tissue sample (left) and immunostaining for myofiber type markers (right) within corresponding areas.

**Figure S2.**
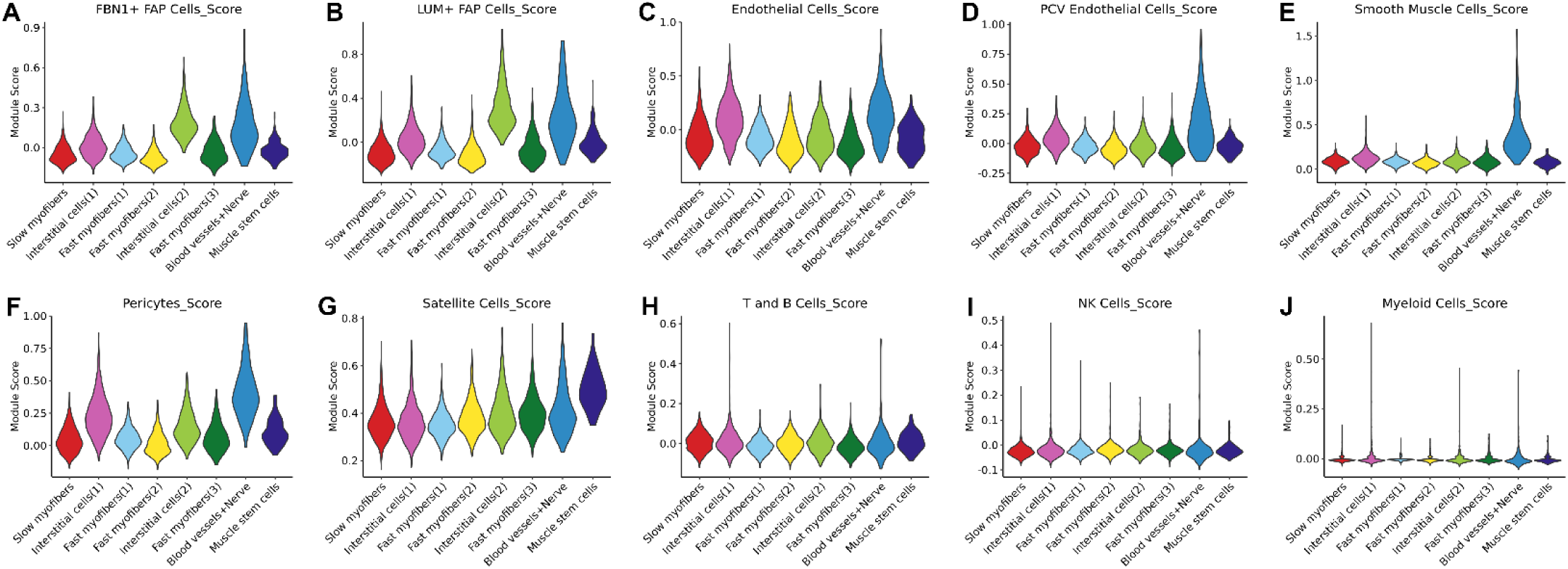
Single cell gene module score integration with spatial transcriptomics clusters. (A-J) Violin plots depicting integration of gene module scores of human muscle single cell populations from the Rubenstein et al.^10^ data with each spatial transcriptomics cluster.

**Table S1**. Number of spots analyzed per sample, with mean transcripts and genes detected per spot. Related to Figure 1.

**Table S2**. Gene expression signatures for each of the 9 clusters identified by spatial transcriptome profiling. Related to Figure 1.

**Table S3**. Differentially expressed genes between sedentary and exercised spots within IHC-defined fast and slow myofiber clusters. Related to Figure 4.

**Table S4**. Gene ontology analysis of biological processes differentially regulated between interstitial cell clusters. Related to Figure 5.

